# Developmental regulation of fear memories by an obesogenic high-saturated fat/high-sugar diet

**DOI:** 10.1101/748079

**Authors:** Julio David Vega-Torres, Arsenio L. Reyes-Rivera, Johnny D. Figueroa

## Abstract

**Background:** Anxiety and stress-related disorders are strongly linked with obesity and the consumption of obesogenic diets. Paralleling clinical findings, we showed that the consumption of an obesogenic diet during adolescence disrupts the structural integrity of amygdalar and prefrontal cortex circuits underlying emotional responses to stress. These abnormalities were associated with a PTSD-like phenotype, including heightened stress reactivity to predator odor trauma, anxiety-like behaviors, and profound learning deficits. The present follow-up study investigates how an obesogenic diet alters aversion-related associative memories across adolescence.

**Methods:** Adolescent Lewis rats were fed for eight weeks with an experimental Western-like high-saturated fat/high-sugar diet (*WD*, 41% kcal from fat) or a matched control diet (*CD*, 13% kcal from fat). Acoustic fear-potentiated startle (FPS) responses were assessed longitudinally at weeks 1, 4, and 8 after commencing the diets to determine the effects of the WD on cued fear conditioning, fear extinction learning, and fear extinction retention.

**Results:** We found that the rats that consumed the WD exhibited substantial attenuation of fear extinction and fear extinction retention when remote memory was tested. One-week WD consumption was sufficient to induce impairments in fear extinction learning. This phenotype was associated with reduced dopamine receptor 1 mRNA levels in the prefrontal cortex. Interestingly, our reconditioning paradigm revealed that early-acquired fear memories were resistant to the disruptive effects of chronic WD consumption on cued fear learning.

**Conclusions:** Our findings demonstrate that consumption of an obesogenic WD during adolescence heightens behavioral vulnerabilities associated with risk for anxiety and stress-related disorders. Given that fear extinction promotes resilience and that fear extinction principles are the foundation of psychological treatments for PTSD, understanding how obesity and obesogenic diets affect the acquisition and expression of fear extinction memories is of tremendous clinical relevance.

**HIGHLIGHTS:**

- Acute WD consumption impairs cued fear extinction learning in a fear-potentiated startle paradigm.
- WD consumption attenuates fear extinction memory retention
- WD consumption during adolescence increases acoustic startle responsivity over time
- Chronic WD consumption decreases dopamine receptor D1 mRNA levels in the prefrontal cortex.

## 1. INTRODUCTION

Early-life trauma is increasingly linked to obesity and the consumption of obesogenic diets (Duncan et al., 2015; Ehlert, 2013; Farr et al., 2015; Masodkar et al., 2016; Mason et al., 2017; Pagoto et al., 2012; Perkonigg et al., 2009; Roenholt et al., 2012; Wolf et al., 2017). The high co-morbidity between obesity and post-traumatic stress disorders (PTSD) suggest that adaptations to trauma may increase the risk for the consumption of obesogenic diets as a result of the traumatic experience (Godfrey et al., 2018; Kalyan-Masih et al., 2016; Michopoulos et al., 2016). There is mounting evidence that exposure to obesogenic diets rich in saturated fat diets and sugars has a direct adverse effect on emotional regulation, anxiety-like behaviors, and neural substrates implicated with stress (Baker and Reichelt, 2016; Boitard et al., 2015; Kalyan-Masih et al., 2016; Ortolani et al., 2011; Reichelt et al., 2015; Sivanathan et al., 2015; Vega-Torres et al., 2018). Therefore, it is possible that early-life exposure to obesogenic diets may predispose individuals to maladaptive stress responses, resulting in increased PTSD risk.

Several lines of evidence suggest that alterations in attention, memory, and learning contribute to the etiology and maintenance of PTSD symptoms (Liberzon and Abelson, 2016; Lissek and van Meurs, 2015). Interestingly, while fear learning emerges early in life, fear memories undergo dynamic changes during adolescence (Baker and Richardson, 2015; Ganella et al., 2017; King et al., 2014). Studies indicate that extinction learning is blunted during adolescence (Pattwell et al., 2012), which has important implications for PTSD treatment. Fear-potentiated startle (FPS) represents a proven and reliable method for examining conditioned fear responses (M. Davis et al., 1993). This method shows notable face validity, construct validity, and predictive validity in the assessment of behaviors and circuits implicated in PTSD. The highly conserved corticolimbic circuit is critical for cue-elicited fear responses and safety learning (Likhtik and Paz, 2015; Likhtik et al., 2014). In particular, the corticolimbic pathway connecting the medial prefrontal cortex (mPFC) and the basolateral complex of the amygdala (BLA), which undergoes dramatic structural reorganization during adolescence (Arruda-Carvalho et al., 2017; Casey, 2015; Silvers et al., 2017) and remains the focus of our recent investigations. While previous studies from our laboratory indicate that this fear circuit is highly vulnerable to the disruptive effects of obesogenic diets (Vega-Torres et al., 2018), to our knowledge, no studies have investigated the effect of obesogenic diets on fear extinction across adolescence.

This follow-up study tested several hypotheses. First, we predicted impaired fear learning in adolescent rats that consumed the obesogenic diet for one week. Not only would this represent a replication of our previous report in adult rats (Vega-Torres et al., 2018), but it would also extend that finding to diet effects independent of obesity-related processes. Second, we predicted attenuated extinction learning and extinction recall in the adolescent rats that consumed the obesogenic diet. Third, we anticipated normal fear associations if the rats were conditioned before the establishment of an obesogenic phenotype. This finding would suggest a time-dependent course of chronic diet effects on the fear neurocircuitry, confirming previous studies (Kalyan-Masih et al., 2016; Vega-Torres et al., 2018). Given the robust effects of obesogenic diets on the dopamine system (Reichelt, 2016; Reyes, 2012), and the modulatory actions of this neurotransmitter on mPFC-amygdala circuit function and FPS responses (Fadok et al., 2009; Onozawa et al., 2011), we hypothesized that the obesogenic diet would reduce dopamine receptor expression in the mPFC and amygdala.

This study demonstrates that associative learning and its inhibition are highly susceptible to the consumption of obesogenic diets, even in the absence of an obesogenic phenotype. These findings reveal a unique interplay between diet and fear extinction learning during adolescence, which may prove informative for understanding risk factors implicated in PTSD. Further, our studies suggest that obesity and the consumption of obesogenic diets may represent a mediator of differential PTSD psychotherapy treatment outcomes.

## 2. METHODS

### Animals

All the experiments were performed following protocols approved by the Institutional Animal Care and Use Committee (IACUC) at the Loma Linda University School of Medicine. A total of 122 rats were used to reach the conclusions of this study (FPS protocol optimization: *n* = 18 rats; cross-sectional experiments: *n* = 72 rats; longitudinal experiments: *n* = 32 rats). Adolescent male Lewis rats (postnatal day, PND 21) were purchased from Charles River Laboratories (Portage, MI, USA). The rationale for the use of Lewis rats is based on the relevant vulnerabilities of this strain to post-traumatic stress (Kalyan-Masih et al., 2016; Vega-Torres et al., 2018). Immediately upon arrival, the rats were housed in groups (2 per cage) with free access to food and water. Dietary manipulations commenced at PND 28. Animals were kept in customary housing conditions (21 ± 2 C, relative humidity of 45%, and 12-hour light/dark cycle with lights on at 7:00 AM). The body weights were recorded once a week or daily during the week of behavioral testing. Food consumption was quantified at least four times per week. The rats were never food or water restricted.

### Diets

The standard chow diet (**CHOW**, 4-gm% fat, product #: 7001) was obtained from Teklad Diets (Madison, WI, USA), while the matched low-fat purified control diet (**CD**, 5-gm% fat, product #: F7463) and Western-like high-saturated fat/high-sugar diet (**WD**, 20-gm% fat, product #: F7462) were obtained from Bio-Serv (Frenchtown, NJ, USA). The macronutrient composition and fatty acid profiles are detailed in **Table 1**. There is an increasing awareness of the impact of diet changes on stress reactivity (Sharma et al., 2013; Vega-Torres et al., 2018) and a need for using adequate matched diets in nutrition research (Pellizzon and Ricci, 2018). Thus, we decided to incorporate a matched low-fat control diet group along with the standard chow diet group. This facilitated the interpretation of the acute diet effects on behavior by diminishing potential confounding factors associated with switching from a CHOW diet to the experimental WD. Given that initially both the CHOW and CD groups had identical biometric and behavioral outcomes, we opted to use the more appropriate CD group as control for the WD group.

**Table 1.** Detailed diet compositions of the CHOW, CD and WD. Dietary fat percentage was ~4% of dry weight in the CHOW, ~5% in the CD, and ~20% in the WD. The WD contained higher levels of total saturated fatty acids (9-fold difference) and total monounsaturated fatty acids (3-fold difference) when compared to the CD. The levels of total polyunsaturated fatty acids were similar between CD and WD. *NR*, not reported by manufacturer or not determined by the mass-spec analyses.

|  | **CHOW** | **CD** | **WD** |
| --- | --- | --- | --- |
| % kcal Carbohydrates | 53 | 68 | 43 |
| % kcal Protein | 34 | 19 | 16 |
| **% kcal Fat** | 13 | 13 | 41 |
| Total kcal/gram | 3.0 | 3.7 | 4.6 |
| **Fatty Acids (g/kg)** |  |  |  |
| C4:0 (Butyric Acid) | NR | 0.72 | 4.89 |
| C6:0 (Caproic Acid) | NR | 0.51 | 3.37 |
| C8:0 (Caprylic Acid) | NR | 0.32 | 2.11 |
| C10:0 (Capric Acid) | NR | 0.75 | 5.08 |
| C12:0 (Lauric Acid) | NR | 0.90 | 5.93 |
| C14:0 (Myristic Acid) | NR | 2.87 | 18.8 |
| C15:0 (Pentadecylic Acid) | NR | 0.31 | 2.01 |
| **C16:0 (Palmitic Acid)** | 7.0 | 10.9 | 56.1 |
| C17:0 (Margaric Acid) | NR | 0.17 | 1.01 |
| C18:0 (Stearic Acid) | 3.0 | 3.34 | 19.6 |
| C20:0 (Arachidic Acid) | NR | 0.11 | 0.29 |
| **Total SFA** | **11.0** | **19.6** | **113** |
| C14:1 (Myristoleate Acid) | NR | 0.22 | 1.46 |
| C16:1 (Palmitoneic Acid) | NR | 0.40 | 2.62 |
| C18:1 (Oleic Acid) | 12.0 | 10.4 | 36.7 |
| **Total MUFA** | **14.0** | **11.6** | **45.5** |
| C18:2<sub>n6</sub> (Linoleic Acid) | 8.0 | 10.9 | 10.2 |
| C18:3<sub>n3</sub> (Alpha-Linoleic Acid) | 1.0 | 0.31 | 0.82 |
| C20:3$n$6 (Dihomo-<br>gamma-linoleic Acid) | NR | <0.06 | 0.22 |
| C20:4$n$6 (Arachinoid<br>Acid) | NR | <0.06 | 0.29 |
| C20:5$n$3<br>(Eicosapentaenoic Acid) | NR | <0.06 | <0.14 |
| C22:6$n$3<br>(Docosahexaenoic Acid) | NR | <0.06 | <0.14 |
| **Total PUFA** | **9.0** | **10.7** | **11.1** |

### Study Design

Behavior testing sessions involved a 20-30 min acclimation period to the testing facility. Following room acclimation, the rats were placed for 5 min inside the acoustic startle reflex (ASR) enclosure and testing chamber and then, returned to their cages. The next day, we measured baseline ASR responses, which were used to generate balanced experimental groups. The ASR-based group matching resulted in an even distribution of rats with similar startle responses in all groups. We conducted *a priori* F power analyses (Cohen, 1992) (repeated measures two-way ANOVA; 3 groups, 2 measurements) using *G*Power* (Faul et al., 2007). Analyses revealed that a minimum of 6 rats per diet group would be sufficient to detect medium effect sizes (*d* = .40) on FPS responses with power (1 - β) set at .80, and α = .05. This follow-up longitudinal study consisted of the following diet groups: CHOW (*n* = 12), CD (*n* = 10), and WD (*n* = 10). Rats were subdivided in two groups: **1)** reconditioned group (RC), rats that were subjected to fear conditioning at PND 35 and re-exposed to the same conditioning protocol at PND 84 (*n* = 6/group), and **2)** late conditioned group (LC), rats that were fear conditioned only at PND 84 (n = 4-6/group). Only data from reconditioned rats is presented in this manuscript. The rats were allowed to consume the diets until completion of the study (from PND 28 to PND 88). The fear-potentiated startle (FPS) paradigm was performed to assess the effect of the diet on cued fear conditioning and fear extinction learning at PND 35 (1-week diet intake = acute effects). We measured ASR responses and fear extinction retention at PND 56 (4 weeks on diet = subchronic effects). ASR and FPS responses were re-assessed at PND 84 (8 weeks diet intake = chronic effects). All the rats were euthanized 24 h following the last FPS protocol. **Figure 1** summarizes the timeline of experimental procedures and behavioral tests.

**Figure 1.**
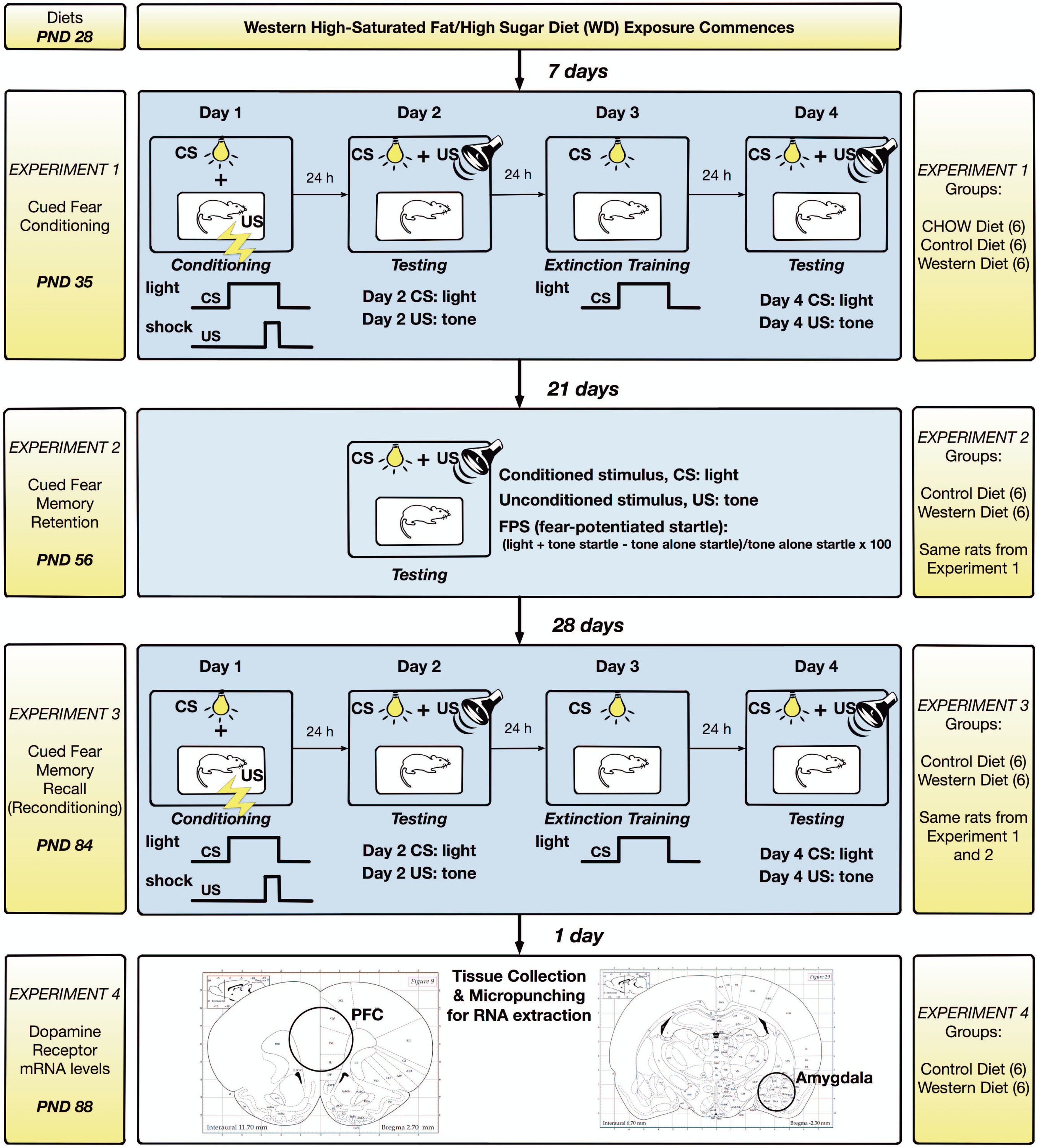
Timeline of experimental procedures. Rats were matched based on their acoustic startle reflex (ASR) responses and allocated to one of three diets: lab chow diet (CHOW), control diet (CD), or Western high-saturated fat diet (WD). The rats were allowed to consume the diets until completion of the study (from PND 28 to PND 88). Please note that the same rats were used throughout the study (*n* = 6 rats per group). *Experiment 1*: A 4-day fear-potentiated startle (FPS) paradigm was used to determine the effects of the WD on cued fear conditioning and fear extinction learning at PND 35. The rats were conditioned to pair a light stimulus (conditioned stimulus, CS) with a foot shock (unconditioned stimulus, US). The conditioning session consisted of 10 CS + US presentations (the light stimulus was paired with a co-terminating foot shock to probe amygdala-dependent learning). Acquisition of the fear memory was assessed 24 h later. One day following fear conditioning testing, the rats were exposed to an extinction-training session. The extinction training session consisted of 30 presentations of the CS alone (light without shock or noise bursts). One day after fear extinction training, we evaluated the acquisition of fear extinction memory using a testing session identical to that used to measure the acquisition of fear. *Experiment 2*: Unconditioned and conditioned responses were re-evaluated at PND 56. *Experiment 3*: The rats were exposed to a second FPS paradigm at PND 84 to reinstate fear. *Experiment 4*: All the rats were euthanized one day after the completion of the FPS protocol (and dietary manipulations) and brains dissected for RNA extraction. Please refer to the study design in the methods section for more specific technical details on procedures and behavioral tests performed in this study.

#### Experiments 1 and 3

The acoustic startle reflex (ASR) and its potentiation (FPS) were evaluated at weeks 1, 4, and 8 following the dietary manipulations. The rats were fear conditioned at week 1 and reconditioned at 8 following the dietary manipulations.

### Acoustic startle reflex (ASR)

The ASR experiments were performed using the SR-Lab acoustic chambers (San Diego Instruments, San Diego, CA, USA). ASR magnitudes were measured by placing animals startle enclosures with sensors (piezoelectric transducers) that convert small movements to voltage. Thus, the magnitude of the change in voltage represents the size of the ASR. Acoustic stimuli intensities and response sensitivities were calibrated before commencing the experiments. The ASR protocol has been previously described by our group (Kalyan-Masih et al., 2016; Vega-Torres et al., 2018). Briefly, experimental sessions were 22 min long and started with a 5 min habituation period (background noise = 55 decibels, dB). The rats were then presented with a series of 30 tones (10 tones at each intensity: 90 dB, 95 dB, and 105 dB) using a 30 sec inter-trial interval (ITI). The acoustic stimuli had a 20 millisecond (ms) duration and the trials were presented in a quasi-random order. Subsequently, the rats were returned to their home cages. Enclosures were cleaned with soap and water and thoroughly dried following each session. Averaged ASR magnitudes were normalized by weight to eliminate confounding factors associated with body weight (Weight-corrected ASR = ASR magnitude in mV divided by body weight at testing day) (Elkin et al., 2006; Gogos et al., 1999; Grimsley et al., 2015; Kalyan-Masih et al., 2016; Vega-Torres et al., 2018). ASR responses were measured at the start of the study (baseline) and 48 h before FPS fear extinction retention testing and reconditioning.

### Fear potentiated startle (FPS)

The fear potentiated startle (FPS) protocol was adapted from Dr. Michael Davis (M. Davis, 2001) and detailed in our previous studies (Vega-Torres et al., 2018). Each FPS session started with a 5 min acclimation period (background noise = 55 dB). During the first session of the paradigm (fear training), the rats were trained to associate a light stimulus (conditioned stimulus, CS) with a 0.6 mA foot shock (unconditioned stimulus, US). The conditioning session involved 10 CS + US pairing presentations. During each CS + US presentation, the light (3200 ms duration) was paired with a co-terminating foot shock (500 ms duration). Light-shock pairings were presented in a quasi-random manner (ITI = 3-5 min). Cued fear acquisition was measured 24 h later. During the second session (fear learning testing), the rats were first presented with 15 startle-inducing tones (*leaders*; 5 each at 90 dB, 95 dB, and 105 dB) delivered alone at 30 sec ITI. Subsequently, the rats were presented with 60 test trials. For half of these test trials, a 20 ms tone was presented alone (**tone alone trials**; 10 trials for each tone: 90 dB, 95 dB, and 105 dB). For the other half, the tone was paired with a 3200 ms light (**light + tone trials**; 10 trials for each pairing: 90 dB, 95 dB, and 105 dB). To conclude the testing session, the rats were presented with 15 startle-inducing tones (*trailers*; 5 each at 90 dB, 95 dB, and 105 dB) delivered at 30 sec ITI. Trials in this session were presented in a quasi-random order (ITI = 30 sec). The startle-eliciting tones had a 20 ms duration. One day after fear conditioning testing, the rats were exposed to a single extinction-training session. The extinction training session consisted of 30 CS alone presentations (light without shock or noise bursts) with a duration of 3700 ms (ITI = 30 sec). One day after fear extinction training, we determined fear extinction acquisition using the same FPS session that was used to measure fear acquisition. It is noteworthy that in this study we shortened the fear extinction training protocol to a single session as opposed to 3 sessions in our previous report in adult rats (Vega-Torres et al., 2018). Unpublished findings from our lab have confirmed that a single extinction training session enables accurate and precise fear extinction measurements without flooring effects in adolescent Lewis rats. We investigated fear learning, fear extinction learning, and fear memory retention by comparing the startle amplitude from light + tone trials (conditioned + unconditioned stimulus, CS + US) relative to tone alone trials (unconditioned stimulus, US). Fear extinction memory retention was assessed using the same testing session that was used to measure fear learning and extinction. FPS data were reported as the proportional change between US and CS + US [%FPS = ((Light + Tone Startle) − (Tone Alone Startle)) / (Tone Alone Startle) × 100] (Walker and M. Davis, 2002). Fear extinction was scored as the FPS difference between the fear learning and the fear extinction testing sessions (pre-extinction FPS – post-extinction FPS). Longitudinal responsivity to leaders and trailers tone alone startle-inducing trials was expressed as percent change from control diet at baseline.

#### Experiment 2

Fear memories were assessed at week 4 following the dietary manipulations. The rats were exposed to the same ASR protocol and FPS *testing session* described in experiments 1 and 3. Please note that the rats were not exposed to the foot shocks in this experiment.

### Experiment 4

#### Real-time quantitative polymerase chain reaction (real-time qPCR)

Brain tissue was collected 24 h following the extinction testing session (after 8 weeks on the dietary manipulations). The rats were euthanized with Euthasol (Virbac, Fort Worth, TX, USA) and perfused transcardially with phosphate buffer saline (PBS). Following the perfusion procedure, the prefrontal cortex and the amygdala were isolated. Total RNA was extracted using Trizol (Invitrogen Life Technologies, Carlsbad, CA). The only change to the recommended protocol was using 1–bromo–3–chloropropane (BCP, 0.1 mL per 1 mL of Trizol) instead of chloroform. BCP was obtained from Molecular Research Center (Cincinnati, OH). RNA concentration was determined on a NanoDrop spectrophotometer (Thermo Scientific, Waltham, MA). We used 1 microgram of the total RNA for cDNA synthesis (iScript cDNA Synthesis Kit, Cat. #170-8891, Bio-Rad Laboratories, Hercules, California, USA). cDNA synthesis protocol was performed according to manufacturer’s instructions. The total volume of the cDNA synthesis reaction mixture was 20 microliters (4 microliters, iScript reaction mix; 1 microliter, iScript reverse trancriptase; 15 microliters, nuclease-free water and 1 microgram of RNA). After completion of cDNA synthesis, 80 microliters of nuclease-free water were added to dilute the 20 microliters of synthesized cDNA. The cDNA was amplified by PCR using the following primer sets: dopamine receptor 1 (DR1, forward: 5’-ATC GTC ACT TAC ACC AGT ATC TAC AGG A-3’; reverse: 5’-GTG GTC TGG CAG TTC TTG GC-3’); and dopamine receptor 2 (DR2, forward: 5’-AGA CGA TGA GCC GCA GAA AG-3’; reverse: 5’-GCA GCC AGC AGA TGA TGA AC-3’). Glyceraldehyde 3-phosphate dehydrogenase (GAPDH, forward: 5’-AGT TCA ACG GCA CAG TCA AG-3’; reverse: 5’-GTG GTG AAG ACG CCA GTA GA-3’) served as housekeeping gene for normalization. Real-time PCR amplification and analyses were carried out on the CFX96 Real-Time PCR Detection System (Bio-Rad Laboratories, Hercules, California, USA). Real-time qPCR conditions were optimized and 25 microliters reactions were prepared. The PCR reactions contained: 12.5 microliters of iQ SYBR Green Supermix (Cat. #170-8882, Bio-Rad Laboratories, Hercules, California, USA), 1 microliter of a mixture of 10 micromolar forward/reverse primer, 6.5 microliters of water, and 5 microliters of the previously synthesized cDNA. The PCR protocol started with 5 minutes at 95°C. This was followed by 40 cycles of: 15 seconds at 95°C for denaturation and 1 minute at 60°C for annealing/extension. The relative levels of DR1 and DR2 mRNA were calculated using the comparative C*t* (crossing threshold). Each sample was normalized to its GAPDH mRNA content. Relative gene expression levels were normalized to CD group and expressed as percent change from control.

### Statistical Analysis

We analyzed the data using Graphpad Prism version 8.0. Shapiro-Wilk statistical analyses revealed that our data sets passed the normality test (*p* > 0.05), indicating that we cannot reject the null hypothesis that the startle responses come from a population which exhibits a normal distribution. The Brown-Forsythe test was used to test for the equality of group variances. When appropriate, the Student’s t-test, one-way ANOVA, or two-way repeated-measures ANOVA were used to examine the effect of diet on gene expression, foot shock reactivity between diet groups, and the influence of diet type, stimulus type, time and interactions on fear responses. Multiple comparisons were made using Sidak’s post-hoc test. The Grubbs’ method was used to investigate outliers. We considered differences significant if *p* < .05. The data is shown as the mean ± standard error of the mean (S.E.M.). We also conducted a post-hoc power analysis with the *G*Power* program (Faul et al., 2007). For repeated measures two-way ANOVA analyses, the statistical power (1-β) for data including the three diet groups was .20 for detecting a small size effect (*d* = .2), whereas the power exceeded .99 for the detection of a large effect size (*d* = .8). The statistical power for all the remaining datasets comparing CD and WD was .24 for detecting a small size effect (*d* = .2), whereas the power was once again more than .99 for the detection of a large effect size (*d* = .8). This indicates that this study is adequately powered at or above a moderate size level (*d* = .4). Therefore, if chosen at random, the probability that rats that consumed the WD will exhibit alterations in fear-related behaviors relative to controls is .61.

## 4. RESULTS

### Bodyweight and food consumption

To determine whether the WD significantly altered body weight and caloric intake relative to the control diets, we measured body mass (weekly and on behavioral testing days) and food consumption (every other day). In agreement with previous findings, we found that the rats that consumed the obesogenic WD showed increased body mass and food consumption. Repeated measures two-way ANOVA analyses revealed significant diet [*F*_(2, 29)_ = 3.56, *p* = .04], time [*F*_(8, 232)_ = 9,216, *p* < .0001], and interaction [*F*_(16, 232)_ = 35.73, *p* < .0001] effects on body weight (**Figure 2A**). Sidak’s post-hoc analyses showed that differences in body weight were statistically significant at week 6 when comparing CHOW and WD groups (5% increase; *p* = .004). This difference in body weight continued increasing and was sustained until completion of study (for weeks 7-9: *p* < .0001). We found significant alterations in body weight that emerged at week 7 when comparing CD and WD groups (6% increase*; p* = .0008), an effect that was sustained until completion of study (weeks 8-9: *p* < .0001). The CD group gained more weight than the CHOW group on week 9 (*p* = .01). Figure 2B shows group body weights during behavioral testing weeks. One-way ANOVA analyses revealed no differences between the diet groups at week 1 [*F*_(2, 29)_ = 1.65, *p* = .21] and week 4 [*F*_(2, 29)_ = 1.26, *p* = .30]. Analyses revealed significant differences in body weights at week 8 [*F*_(2, 29)_ = 13.22, *p* < .0001]. Sidak’s post hoc analyses showed increased body weight in the WD group (8% increase relative to CHOW rats, *p* = .0001; 7% increase relative to CD rats, *p* = .001). Similarly, we found significant diet [*F*_(2, 13)_ = 95.57, *p* < .0001], time [*F*_(8, 104)_ = 69.23, *p* < .0001], and interaction [*F*_(16, 104)_ = 2.95, *p* = .0005] effects on food consumption (**Figure 2C**). Analyses revealed a significant 19% increase in caloric intake in WD rats relative to CHOW rats after the first week on the diets. This striking difference in food intake was persistent and reached more than 25% difference in food consumption (for weeks 1-9: *p* < .01). Comparably, we found increased caloric intake in WD rats relative to CD rats (*p* < .01). The differences in food consumption between these groups emerged at week 2 (9% increase) and extended until the completion of study (up to 26%; for weeks 2-9: *p* < .01).

**Figure 2.**
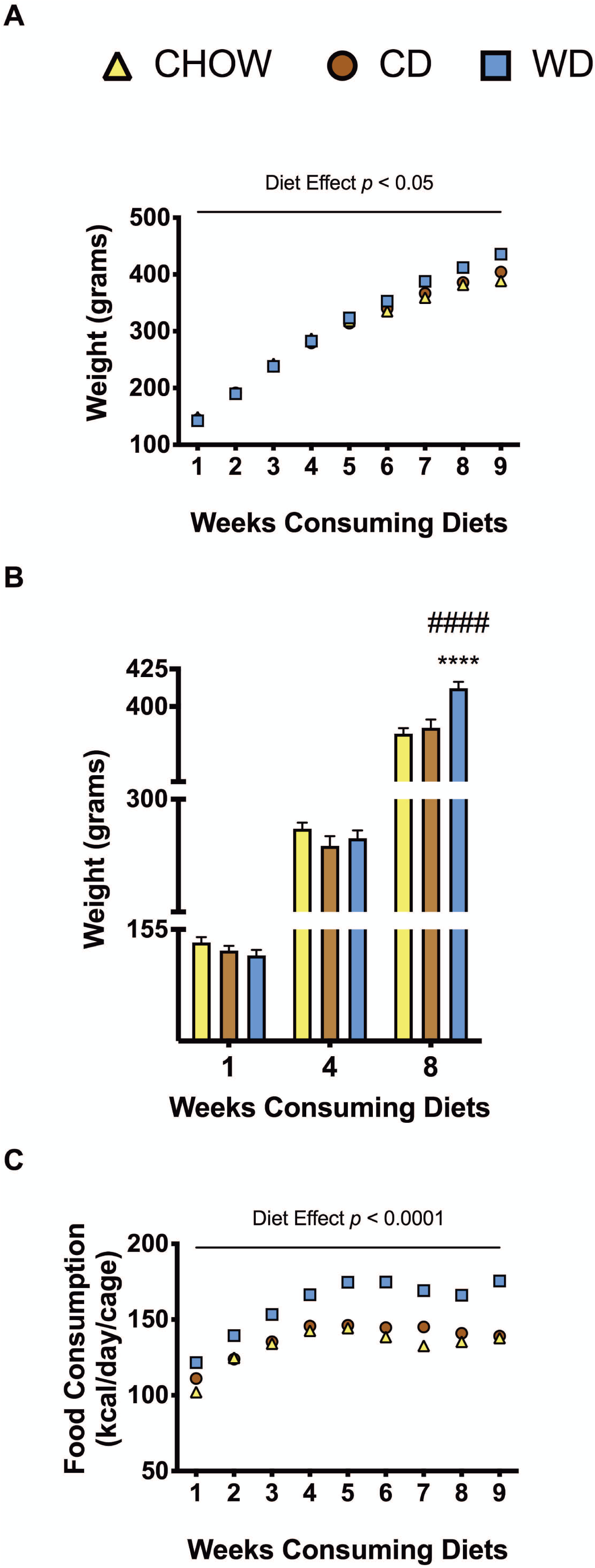
Body mass and caloric intake changes. **(A)** Average weekly body weight in grams for CHOW, CD and WD groups. Differences in body mass emerged at week 6 (5% increase relative to CHOW, *p* = .004) and at week 7 (6% increase relative to CD*, p* = .0008) *n* = 10-12 rats/group. **(B)** Body weights during behavioral testing weeks. The WD group had a significant increase in body weight at week 8 when compared to both CHOW (8% increase; p = .0001) and CD (7% increase; *p* = .001). **(C)** Average daily caloric intake in kilocalories per cage. WD group consumed more calories than CHOW (for weeks 1-9: *p* < .01) and CD group (for weeks 2-9: *p* < .01); n = 5/6 cages/group. Error bars are S.E.M.

### Experiment 1: Acute WD exposure attenuates fear extinction learning

We recently reported that chronic consumption of an obesogenic WD during adolescence leads to impairments in fear-related associative learning and extinction in adult rats (Vega-Torres et al., 2018). This follow-up study investigated the acute, subchronic, and chronic effects of adolescent WD consumption on cued fear conditioning and fear extinction (**Figure 1**). We used the acoustic startle reflex (ASR) to balance the rats in each study group based on their unconditioned responses to acoustic stimulation. Following ASR matching, the CD and WD rats were exposed to the novel diets and fed *ad libitum* for one week before behavioral testing. CHOW animals were maintained on the standard lab chow. The acute WD effect on fear responses was investigated using the fear-potentiated startle (FPS). Rats were conditioned to learn the aversive association between a light (conditioned stimulus, *CS*) and a foot shock (unconditioned stimulus, *US*) during the first day of the FPS paradigm. We found that the diet type had no significant effect on the behavioral reactivity to the foot shocks (reactivity is defined as the maximun startle magnitude after the foot shock) [*F*_(2, 15)_ = 1.51, *p* = .25] (data not shown). Acquired fear was tested 24 h after conditioning and defined as significant differences in startle amplitudes between the US (tone alone) and the CS + US (light + tone). While CS + US trials increased startle responses relative to US [*F*_(1 15)_ = 67.60, *p* < .0001], our analyses revealed no diet [*F*_(2, 15)_ = 1.30, *p* = .30] or interaction effects on startle responses between trial type [*F*_(2, 15)_ = .97, *p* = .40] (**Figure 3A**). Surprisingly, we found that one-week WD consumption did not alter fear learning. Sidak’s post-hoc showed that all the diet groups were able to learn the aversive light + shock associations (for *US* vs. *CS + US*: CHOW, *p* = .004; CD: *p* < .0001; WD: *p* = .001). Interestingly, we found significant differences in pre-extinction FPS responses between groups [*F*_(2, 15)_ = 3.78, *p* = .04] (**Figure 3B**). Analyses showed increased pre-extinction FPS in CD rats relative to CHOW-fed rats (*p* = .039). No differences in pre-extinction FPS responses were found when comparing CHOW vs. WD (*p* = .252) and CD vs. WD (*p* = .549).

**Figure 3.**
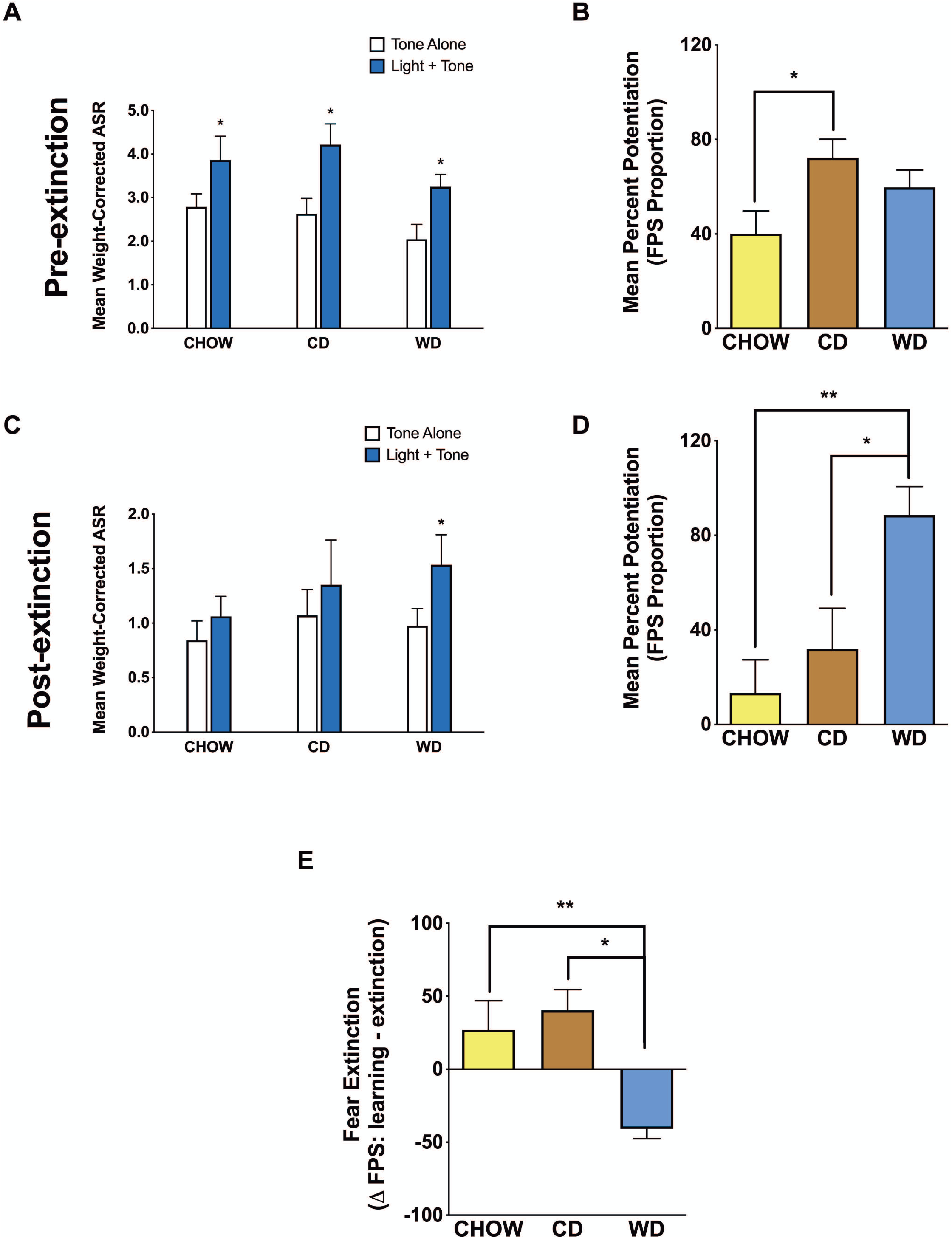
Short-term WD exposure attenuates fear extinction learning in adolescent rats. **(A)** Average weight-corrected ASR responses for the unconditioned (tone alone) and conditioned (light + tone) trials during fear learning testing. All groups showed significant differences between unconditioned and conditioned responses (\*\*\*\**p* < .0001; \*\**p* < .01; *n* = 6 rats/group). **(B)** Fear-potentiated startle (FPS) values from fear learning session (FPS day 2). CD group showed a significant increase in FPS when compared to the CHOW (*p* = .03) (*n* = 6 rats/group). **(C)** Average weight-corrected ASR responses for the unconditioned (tone alone) and conditioned (light + tone) trials during fear extinction testing. Only the WD-fed rats showed significant differences between unconditioned and conditioned responses (*p* = .03; *n* = 6 rats/group). **(D)** Fear-potentiated startle (FPS) values from fear extinction testing session. The WD group showed a significant increase in FPS when compared to CHOW (*p* = .006) and CD (*p* = .03) groups showed a significant reduction in FPS when compared to acquired fear FPS responses (*n* = 6 rats /group). **(E)** Difference in FPS responses from day 2 (pre-extinction FPS) and day 4 (post-extinction FPS) used as an index of fear extinction. WD group showed a significant reduction in fear extinction relative to CHOW (*p* = .02) and CD (*p* = .007) (*n* = 6 rats/group). Error bars are S.E.M.

Fear extinction training and fear extinction testing were performed on days 3 and 4 of the FPS paradigm, respectively. Successful fear extinction is defined as similar startle responses when comparing the US trials to CS + US trials (Vega-Torres et al., 2018). Similar to fear conditioning, the analyses revealed a significant stimulus effect [*F*_(1, 15)_ = 9.56, *p* = .007], revealing differences in startle responses between US and CS + US trials (**Figure 3C**). Diet [*F*_(2, 15)_ = .49, *p* = .62] and interactions between factors [*F*_(2, 15)_ = .84, *p* = .45] did not alter US and CS + US startle responses during post-extinction testing. The WD group exhibited significant impairments in fear extinction learning after only one week on the diet, as evidenced by higher CS + US startle responses relative to startle responsivity to US stimuli (*p* = .03). CHOW (*p* = .63) and CD (*p* = .44) rats showed similar startle responses to the CS + US relative to the US (**Figure 3C**).

ANOVA revealed significant post-extinction FPS differences between the diet groups [*F*_(2, 15)_ = 7.19, *p* = .006] (**Figure 3D**). We found that the WD rats exhibited increased post-extinction FPS relative to the CHOW (*p* = .006) and the CD (*p* = .03) rats. CHOW and CD groups showed similar post-extinction FPS responses (*p* = .64).

The difference in FPS responses from pre-extinction (day 2) to post-extinction (day 4) was used as an index of fear extinction. We found significant fear extinction differences between diet groups [ANOVA *F*_(2, 14)_ = 7.33, *p* = .007] (**Figure 3E**). The rats that consumed the WD exhibited reduced fear extinction relative to CHOW (*p* = .02) and CD (*p* = .007) rats. CHOW and CD rats exhibited similar fear extinction (*p* = .80). On the basis of the similarities in food consumption, body weight, and fear extinction between the CHOW and CD groups, we decided to use the more appropriate matched purified ingredient diet group to control for the experimental WD for the remainder of the study.

### Experiment 2: WD consumption reduces long-term fear memories

Following four weeks on the diets (three weeks after fear extinction testing), we investigated the effects of the WD on long-term extinguished fear responses. We found no significant differences in ASR responsivity between groups (CD: 1.17 ± .14 vs. WD: 1.26 ± .48; *t*_(10)_= .17, *p* = .86) (data not shown). As anticipated, we found larger CS + US startle responses relative the US, even at three weeks after fear extinction training. Analyses revealed a significant stimulus effect [*F*_(1, 10)_ = 9.11, *p* = .01], but no significant diet [*F*_(1, 10)_ = .07, *p* = .79] or interactions between the factors [*F*_(1, 10)_ = .04, *p* = .84] effects (**Figure 4A**). FPS responses were similar between groups at week 4 (CD: 44.73 ± 15.87 vs. WD: 40.95 ± 6.82; *t*_(10)_= .22, *p* = .83) (**Figure 4B**). Interestingly, we found that the WD rats exhibited a larger FPS reduction relative to week 1 (CD: −12.85 ± 18.78 vs. WD: 47.58 ± 12.63; *t*_(10)_= 2.67, *p* = .02) (**Figure 4C**). These findings indicate that although both the CD and WD groups still maintain significant long-term aversive learning-related responses, the WD rats exhibited accelerated fear memory extinction from week 1.

**Figure 4.**
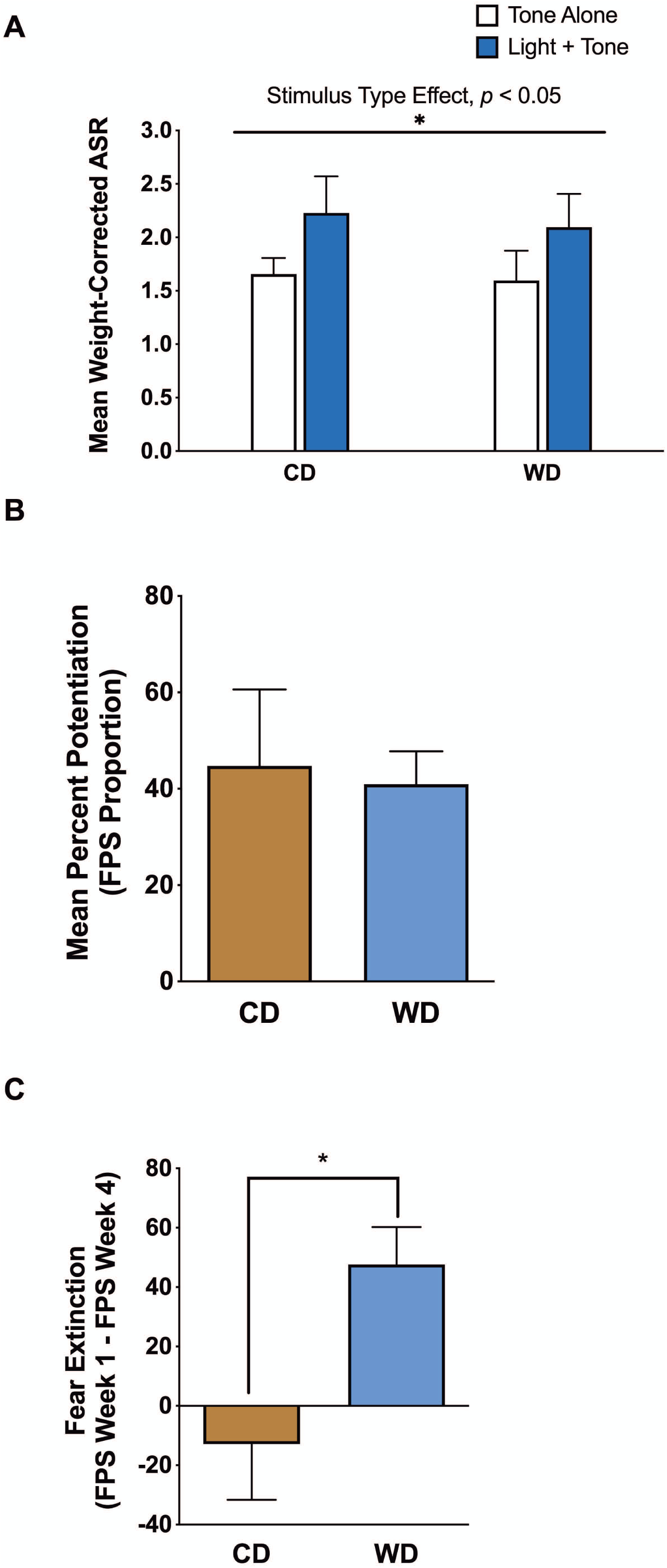
Subchronic WD consumption facilitates spontaneous fear memory extinction. **(A)** Average weight-corrected ASR responses for the unconditioned (tone alone) and conditioned (light + tone) trials during fear extinction testing at three weeks after conditioning. Analyses showed a significant difference between CS + US startle responses relative the US [*F*_(1, 10)_ = 9.10, *p* = .02] (*n* = 6 rats/group). **(B)** Fear-potentiated startle (FPS) values from fear extinction session at three weeks after conditioning. Both diet groups showed similar fear extinction FPS responses (*p* > 0.05; *n* = 6 rats/group). **(C)** Difference in FPS responses from week 1 (post-extinction FPS) to week 4 FPS used as an index of fear extinction. The WD group showed a significant increase in FPS reduction relative to week 1 (p = .02; *n* = 6 rats/group). Error bars are S.E.M.

### Experiment 3: Chronic WD consumption attenuates fear extinction memory retention and delays fear extinction expression

Following 8 weeks on the diets, we performed the ASR protocol to investigate the chronic effect of the WD on startle responses. Analyses showed no significant differences between CD and WD (CD: 1.53 ± .21 vs. WD: 1.55 ± .19; *t*_(10)_= .09, *p* = .93) after 8 weeks on the diets (data not shown). To determine whether the WD affected remote fear memories and vulnerabilities to subsequent stressful events during adulthood, we investigated fear expression after repeated cued fear conditioning. We performed reconditioning in the initial context and under the same experimental conditions to determine whether this reminder procedure would be effective in restoring fear responses and to confirm our studies showing impairments in fear acquisition (Vega-Torres et al., 2018). We found no differences between diet groups on the reactivity to the foot shocks (CD: 9.12 ± .86 vs. WD: 7.69 ± .60; *t*_(10)_= 1.37, *p* = .20) (data not shown).

Subsequently, we confirmed the re-acquisition of the aversive associations. Although we found no effects of the diet [*F*_(1, 10)_ = .60, *p* = .46] or interactions [*F*_(1, 10)_ = .20, *p* = .66], our analyses revealed that the rats were able to acquire the fear associations [*F*_(1, 10)_ = 33.36, *p* = .0002] (**Figure 5A**). Post hoc testing showed that both CD and WD rats were able to exhibit the aversive associations (CD: *p* = .007; WD: *p* = .003). Analyses for pre-extinction FPS responses revealed no significant differences between groups (CD: 70.42 ± 15.11 vs. WD: 80.65 ± 21.06; *t*_(10)_= .39, *p* = .70) (**Figure 5B**).

**Figure 5.**
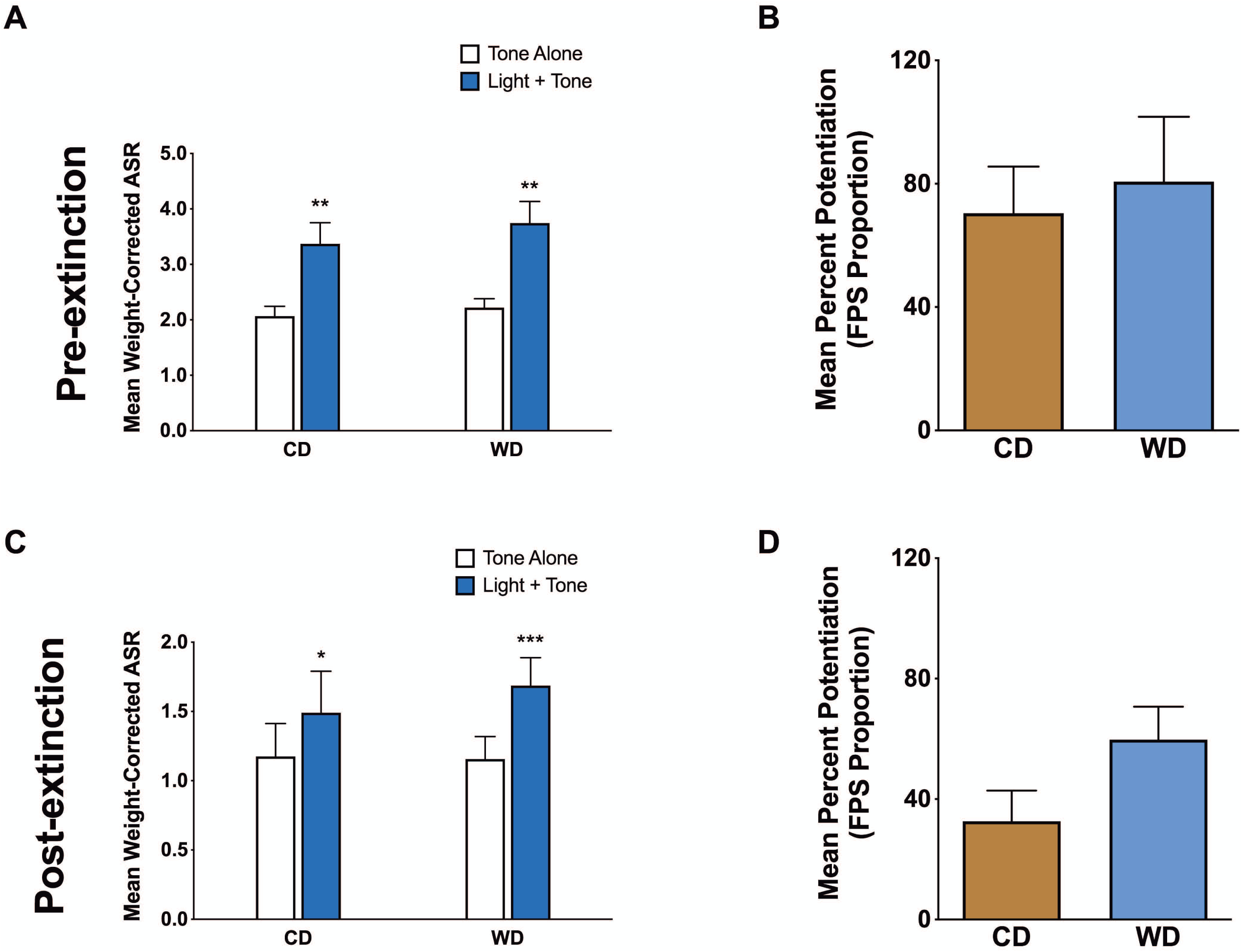
Effect of chronic WD consumption on fear extinction memory. **(A)** Average weight-corrected ASR responses for the unconditioned (tone alone) and conditioned (light + tone) trials during fear learning testing. Both CD and WD showed significant differences between unconditioned and conditioned responses (CD: *p* = .007; WD: *p* = .003; *n* = 6 rats/group). **(B)** Fear-potentiated startle (FPS) values from fear learning session (FPS day 2). Both diet groups showed similar pre-extinction FPS responses (*p* > 0.05; *n* = 6 rats/group). **(C)** Average weight-corrected ASR responses for the unconditioned (tone alone) and conditioned (light + tone) trials during fear extinction testing. Similar to pre-extinction both CD and WD showed significant differences between unconditioned and conditioned responses (CD (p = .02) and WD (p = .0006); *n* = 6 rats/group). **(D)** Fear-potentiated startle (FPS) values from fear extinction testing session. There were no significant effects on post-extinction FPS responses between CD vs. WD (*p* > 0.05; *n* = 6 rats/group). Error bars are S.E.M.

While analyses showed a significant stimulus effect between US and CS + US responses [*F*_(1, 10)_ = 36.78, *p* = .0001], the diet [*F*_(1, 10)_ = .08, *p* = .79] and interactions between factors [*F*_(1, 10)_ = 2.37, *p* = .15] showed no significant effects (**Figure 5C**). Our analyses demonstrated that the US vs. CS + US differences were significant for both CD (*p* = .02) and WD (*p* = .0006). We found a trend towards larger FPS responses in the WD rats relative to CD rats at 8 weeks post-diet consumption (CD: 32.63 ± 10.17 vs. WD: 59.73 ± 10.94; *t*_(10)_= 1.81, *p* = .09) (**Figure 5D**). Diet groups showed similar extinction from learned associations (difference between pre-extinction FPS and post-extinction FPS) (CD: 37.78 ± 15.59 vs. WD: 20.92 ± 21.49; *t*_(10)_= .63, *p* = .53) (data not shown).

In light of findings that show changes in fear and stress reactivity during the transition into and out of adolescence (Chen et al., 2018; Pattwell et al., 2012; 2013; Spear, 2000), we used a neurodevelopmental approach to understand how WD consumption alters the trajectories of fear extinction. By using a conditioning-reconditioning paradigm, we aimed to examine fear responses to immediate extinction as well as long-term fear extinction memories, spontaneous recovery, and fear memory recall. **Figure 6** shows acoustic startle magnitudes of US and CS + US, as rats transitioned from in and out of mid adolescence (CD group is presented in **Figure 6A** and WD group presented in **Figure 6B**). In an attempt to determine whether the WD-induced deficits in fear extinction were associated with delayed acquisition of fear extinction or memory retention, we divided the extinction testing session in two different blocks for each time point (late and early: 18 trials each, 9 US and 9 CS + US). We found that the CD group exhibited similar post-extinction ASR responses to US relative to CS + US across all the blocks and time points studied. Analyses showed a significant main effect of the extinction testing block on ASR responses [*F*_(3, 15)_ = 5.68, *p* = .008]. We found no significant stimulus [*F*_(1, 5)_ = 2.94, *p* = .15] and interaction [*F*_(3, 15)_ = .24, *p* = .87] effects on ASR responses in controls, revealing that this group acquired fear extinction memories and exhibited fear extinction retention. (**Figure 6A**). In contrast, the WD group showed delayed fear extinction retention, as revealed by significant ASR differences between US and CS + US during the early block of extinction testing. We found significant extinction testing block [*F*_(3, 15)_ = 5.64, *p* = .009], stimulus type [*F*_(1, 5)_ = 8.69, *p* = .03] and interaction [*F*_(3, 15)_ = 3.47, *p* = .04] effects on ASR responses for the rats that consumed the WD. Post-hoc analyses revealed that the ASR magnitudes for the US were significantly different from the CS + US-induced ASR responses during the early extinction block, indicating a delayed fear extinction response (week 1, *p* = .008; week 8, *p* = .001) (**Figure 6B**). We found that, relative to CD rats, the WD rats exhibited increased FPS responses during the initial trials of the fear extinction testing session [*F*_(1, 9)_ = 6.70, *p* = .04] (**Figure 6C**). The week of testing [*F*_(1, 9)_ = .75, *p* = .40] and interactions between factors [*F*_(1, 9)_ = .001, *p* = .97] showed no significant influence on FPS responses during the early phase of post-extinction testing.

**Figure 6.**
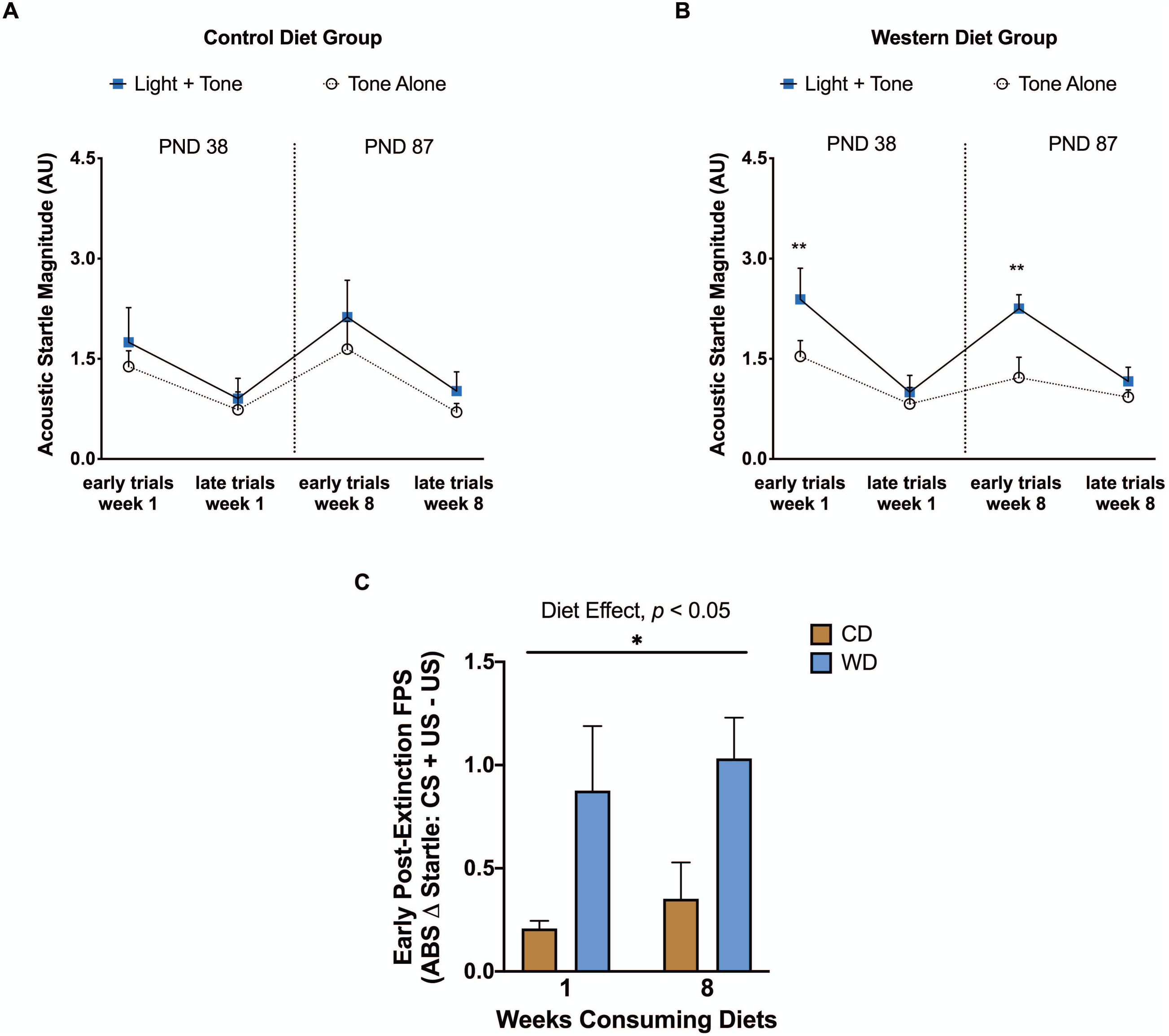
WD consumption leads to fear extinction retention impairments across adolescence. Acoustic startle magnitudes of US and CS + US during fear extinction testing sessions at week 1 and week 8 (two different blocks for each time point; late and early: 18 trials each, 9 US and 9 CS + US). **(A)** CD rats exhibited fear extinction and fear extinction retention (US vs. CS + US: *p* > 0.05; *n* = 6 rats/group). **(B)** WD rats showed marked impairments in fear extinction memory retention. This effect was particularly significant during the early part of the extinction testing session, suggesting delayed fear extinction memory recall or expression (US vs. CS + US: week 1, *p* = .008; week 8, *p* = .001; *n* = 6 rats/group). Error bars are S.E.M.

Given the increased reactivity in post-extinction FPS, we reasoned that these alterations in anxiety-like ASR responses would also be reflected before extinction training, particularly when rats are exposed to the light (CS). Thus, we analyzed startle reactivity to leader (before CS presentation) and trailer (after CS presentation) trials during the pre-extinction session. Longitudinal analyses of the ASR responsivity to the leader trials (block before presentation of light stimuli) showed a significant time point [*F*_(2, 30)_ = 3.49, *p* = .04] effect while no significant diet [*F*_(1, 30)_ = .14, *p* = .70] effect or interaction between the factors [*F*_(2, 30)_ = 1.04, *p* = .36] (**Figure 7A**). Notably, longitudinal analyses of the ASR responsivity to the trailer trials (block after the presentation of light stimuli) showed a significant diet [*F*_(1, 30)_ = 4.72, *p* = .03] and time point [*F*_(2, 30)_ = 44.28, *p* < .0001] effect while no significant interaction between the factors [*F*_(2, 30)_ = 2.32, *p* = .11] (**Figure 7B**). Post hoc revealed a significant diet effect at week 8 (p = .03) (for week 1, *p* = .98; for week 4, *p* = .43). This result suggests increased sensitization, deficits in ASR habituation, and overall, a potential anxiogenic effect of the WD. To confirm the potential impairments in ASR habituation, we also assessed startle responses during the fear extinction testing session by comparing the first 15 tone trials (leaders) and the last 15 tone trials (trailers) of the session (Vega-Torres et al., 2018). We found a significant main effect of the trial type on ASR habituation [trial type: *F*_(1, 10)_ = 10.54, *p* = .009; diet type: *F*_(1, 10)_ = .01, *p* = .92; interactions: *F*_(1, 10)_ = 1.31, *p* = .28]. Analyses showed a significant decrease in the trailer tones relative to the leaders in the CD group (*p* = .02) (data not shown). Similar to previous studies, we found that chronic WD consumption impaired the habituation of the ASR (leader vs. trailer: *p* = .31; data not shown).

**Figure 7.**
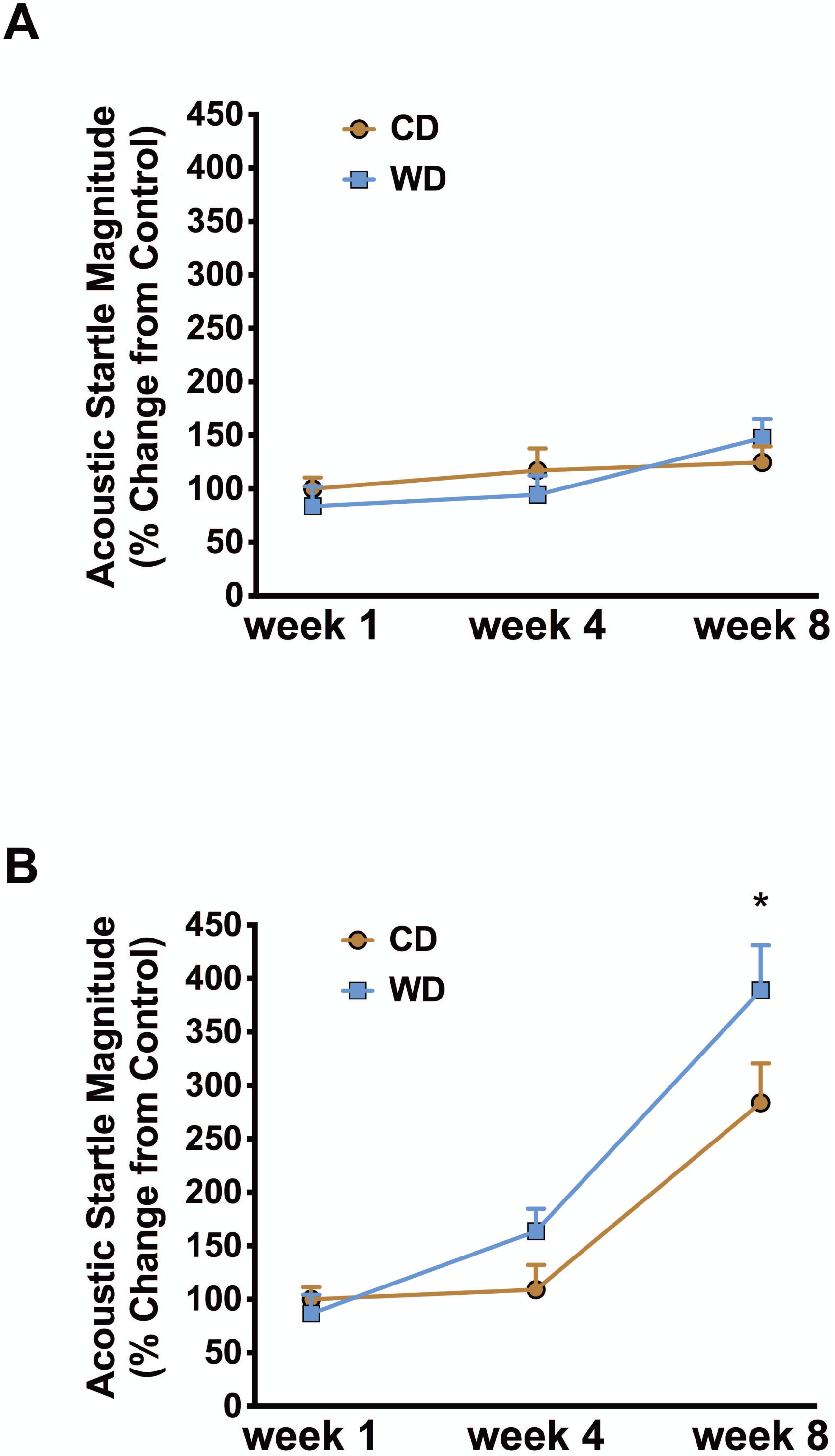
WD consumption alters ASR responses to trailer trials. **(A)** Longitudinal ASR responsivity to leader trials across time expressed as percent change from control at baseline. Both diets showed similar ASR response to leader trials [*F*_(1, 30)_ = .14, *p* = .70; *n* = 6 rats/group]. **(B)** Longitudinal ASR responsivity to trailer trials across time expressed as percent change from control at baseline. WD group showed a significant increase in ASR response to trailers trials at week 8 (*p* = .03) when compared to CD (*n* = 6 rats/group). Error bars are S.E.M.

### *Experiment 4:* Chronic WD consumption attenuates DR1 mRNA levels in the PFC

There is evidence that the activation of dopamine receptors in the prefrontal cortex (PFC) and amygdala are critical for fear learning and extinction (Abraham et al., 2014). Thus, to gain insight into potential molecular targets implicated in the observed fear impairments, we measured the mRNA levels of the dopamine receptor 1 and 2 (DR1 and DR2, respectively) in the PFC and amygdala. Brain tissue was collected 24 h following the extinction testing session. We found that the WD significantly reduced the DR1 mRNA levels in the PFC. Two-way ANOVA analyses showed a significant diet effect [*F*_(1, 10)_ = 8.07, *p* = .01], but no significant receptor type [*F*_(1, 10)_ = .54, *p* = .47] effect or interaction between factors [*F*_(1, 10)_ = 3.59, *p* = .08] (**Figure 8A**). Post hoc revealed significant diet effects in the PFC for DR1 (p = .006) and not for DR2 (p = .66). Unexpectedly, there was no significant diet effect in the amygdala. Analyses showed no significant diet effects [*F*_(1, 9)_ = 1.33, *p* = .27], receptor type effects [*F*_(1, 9)_ = .87, *p* = .37], or interactions between factors [*F*_(1, 9)_ = .0002, *p* = .98] in the amygdala (**Figure 8B**).

**Figure 8.**
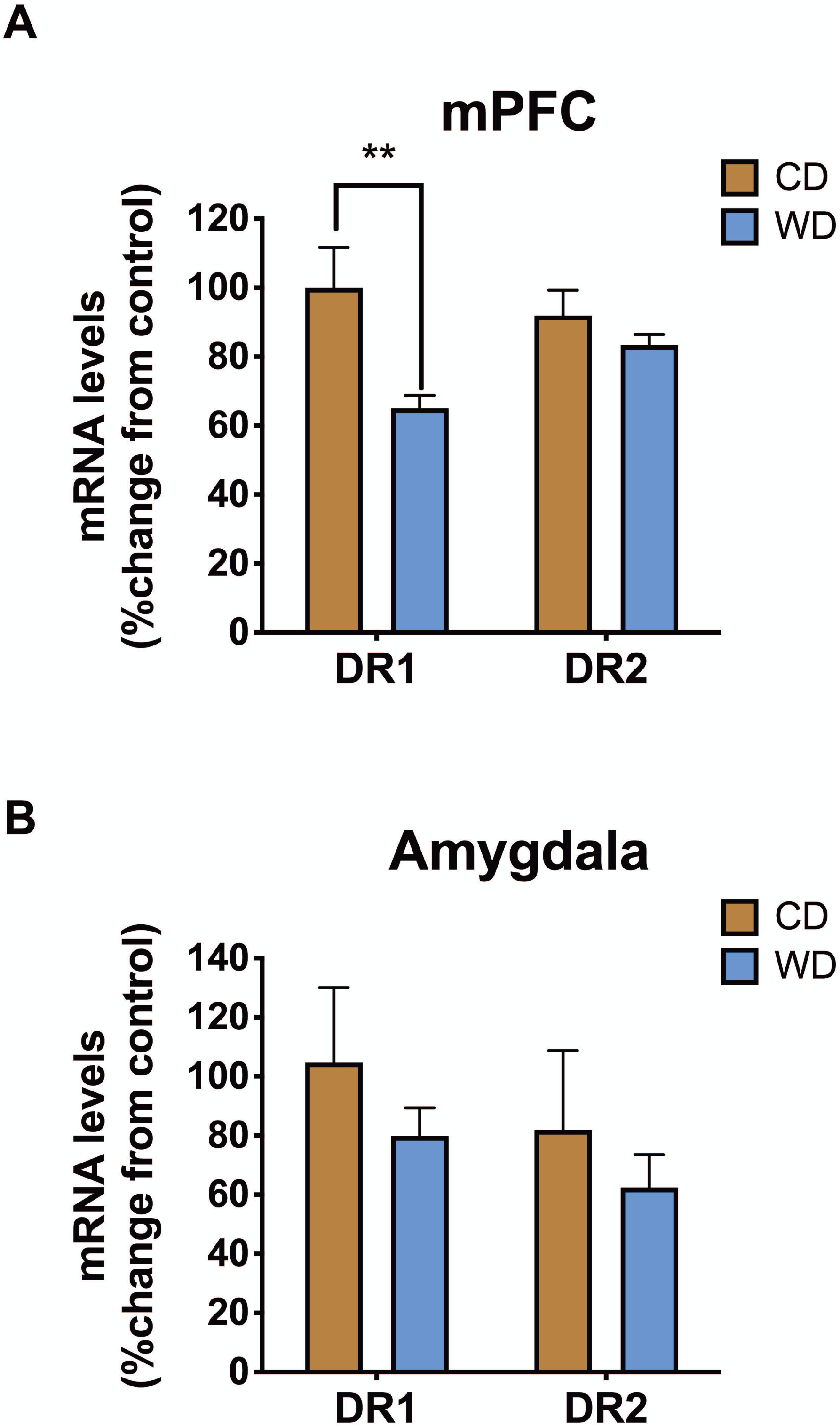
WD consumption reduces dopamine receptor 1 mRNA levels in the prefrontal cortex. Brain tissue was collected 24 h following post-extinction testing. **(A)** qRT-PCR analyses demonstrate reduced dopamine receptor 1 (DR1) mRNA levels in WD rats relative to controls (approximately 35% decrease; posthoc *p* = .006; *n* = 6 rats/group). Diet effects were specific for DR1, as DR2 levels did not change significantly with the dietary manipulation (*p* > 0.05). **(B)** We found no significant differences in DR1 and DR2 mRNA levels in the amygdala (*p* > 0.05; *n* = 5 rats/group). Error bars are S.E.M.

## 5. DISCUSSION

Childhood trauma survivors who suffer from post-traumatic stress disorder (PTSD) are at high risk for developing obesity and metabolic disorders (Assari et al., 2016; Brewerton and ONeil, 2016; Llabre and Hadi, 2009; Ramirez and Milan, 2016; Roenholt et al., 2012). Previous studies support these observations while demonstrating a novel directionality in the relationship between early-life posttraumatic stress reactivity and diet-induced obesity (DIO) (Kalyan-Masih et al., 2016; Vega-Torres et al., 2018). We recently showed that consumption of an obesogenic Western-like high-saturated fat/high-sugar diet (WD) during adolescence heightens stress reactivity while altering key substrates implicated in PTSD (Kalyan-Masih et al., 2016; Vega-Torres et al., 2018). In this follow-up study, we expanded these observations and investigated how WD consumption alters fear responses across time, as rats transitioned into and out of mid-adolescence; a critical period in the development of the fear circuitry. We tested the acute (1 week) and chronic (8 weeks) WD effects, relative to an ingredient matched low-fat semi-purified control diet, on cued fear conditioning, fear extinction, fear extinction recall, and fear reconditioning using the fear-potentiated startle (FPS) paradigm. The primary findings of this study are: **(a)** acute consumption of an obesogenic WD impaired cued fear extinction learning, **(b)** WD consumption resulted in deficits in fear extinction memory retention across multiple time points, **(c)** chronic WD consumption impaired the habituation of the acoustic startle reflex (ASR), **(d)** chronic WD consumption significantly reduced dopamine receptor D1 mRNA levels in the prefrontal cortex, a critical brain structure implicated in regulating conditioned fear responses. Altogether, this follow-up study provides preliminary evidence that behavioral and molecular substrates implicated in fear are selectively vulnerable to the consumption of an obesogenic WD. Despite the possible limitations associated with small sample numbers, our findings also serve to emphasize the caution that must be exercised when interpreting experimental outcomes when unmatched diets are used as controls.

### Short-term WD consumption attenuates fear extinction learning

An important finding of this study is that short-term WD consumption was sufficient to attenuate extinction learning of cued fear. Fear extinction has been characterized as an active form of inhibitory learning that allows for the adaptive regulation of conditioned fear responses (Myers and M. Davis, 2002). The inability to consolidate extinction memory and inhibit conditioned fear under safe conditions underlies some of the hallmark symptoms of anxiety and stress-related disorders (McGuire et al., 2016; Milad et al., 2009; Waters and Pine, 2016). It is now increasingly recognized that cognition, attention, mood, and anxiety disorders have a nutritional component or are promoted by poor dietary habits. Our results are consistent with evidence showing that the consumption of obesogenic diets can attenuate cognitive functions in humans and rodents in as little as 3-7 days (Beilharz et al., 2016; 2014; Edwards et al., 2011; Holloway et al., 2011; Kanoski and Davidson, 2010; Sobesky et al., 2016). The findings of this study indicate that the adverse effects of consuming obesogenic diets expand to additional cognitive domains regulating emotional memories and fear. This is in agreement with new studies showing that adolescent rats that consume high-fat/high-sugar diets exhibit delayed spontaneous extinction and impaired extinction retention of fear-related behaviors (Baker et al., 2016; Reichelt et al., 2015; Vega-Torres et al., 2018). Our findings extend beyond those reported to date by showing delayed fear extinction memory retention and significant fear extinction learning deficits independent of effects associated with obesity and metabolic disturbances.

This study employed a fear conditioning and delayed reconditioning paradigm to identify the longitudinal effects of the WD on post-extinction reconditioning. While our previous studies show that chronic WD consumption attenuates new fear memory acquisition and key neural substrates implicated in fear (Vega-Torres et al., 2018), an intriguing and novel finding of this study is that the WD rats exhibited normal fear conditioning responses if they were conditioned early during adolescence and retested during adulthood. Together, our studies suggest that learned associations acquired before diet-induced structural alterations to the fear neurocircuitry are resistant to the disruptive effects of a WD. In other words, fear memories may be consolidated and stored in accessible long-term memory prior to the damaging effects of the diet. Another possible explanation to our data is that the substrates underlying acquisition of original fear memory are more vulnerable to the disruptive effects of chronic WD consumption, relative to fear memory retrieval and reconsolidation. This idea is supported by studies showing these behavioral processes are mediated by distinct neural projections and mechanisms (An et al., 2018; Johansen et al., 2011; Laurent and Westbrook, 2008; Y. Li et al., 2013; Milad and Quirk, 2002; Seidenbecher et al., 2003). Although evidence that PTSD patients exhibit abnormal fear learning remains controversial, a recent study shows that maltreated children exhibit marked deficits in associative learning during fear conditioning (McLaughlin et al., 2016). Future experiments are required to empirically determine how early-acquired fear memories may be “protected” from the unfavorable effects of a chronic obesogenic diet consumption on fear conditioning.

This study confirms our previous findings showing that rats that consume obesogenic diets may incorrectly asses the level of threat (Kalyan-Masih et al., 2016; Vega-Torres et al., 2018). Evidence supports that impairments in attention and threat discrimination may heighten the risk for anxiety and stress-related psychopathology (Block and Liberzon, 2016; López-Aumatell et al., 2009a; 2009b). Evidence shows that ASR amplitude increases during adolescence, which has been associated with physical development (i.e., increased body weight and muscular tone) (Goepfrich et al., 2017; Pietropaolo and Crusio, 2009; Rybalko et al., 2015) and seems to be dependent on the intensity of the acoustic stimuli (Rybalko et al., 2015; Vega-Torres et al., 2018). The findings of this study are consistent with these reports while showing increased startle responsivity in trailer blocks during ontogeny. This study demonstrates that chronic WD consumption leads to increased ASR responses and deficits in the habituation of the ASR, behavioral proxies for maladaptive stress reactivity and anxiety disorders (Conti and Printz, 2003; Shalev et al., 2000). Taken together, the behavioral outcomes reported here support the notion that perturbations in attention, threat discrimination, and startle habituation may heighten vulnerability for anxiety and stress-related disorders in individuals that consume obesogenic diets.

### Obesogenic diet leads to abnormal maturation of the neural and molecular substrates governing fear extinction

A number of studies indicate that the medial prefrontal cortex (mPFC) and the basolateral complex of the amygdala (BLA) are critical to the acquisition and expression of conditioned fear (M. Davis, 2006; Quirk et al., 2006; Sotres-Bayon et al., 2004). The highly conserved neurocircuitry connecting the mPFC and the amygdala plays a critical role in the extinction of fear memories (Janak and Tye, 2015; Milad and Quirk, 2002; Phelps et al., 2004) and is abnormal in PTSD patients (Gilboa et al., 2004; Koenigs and Grafman, 2009). The PFC and amygdala undergo striking structural changes during adolescence (Jalbrzikowski et al., 2017), providing a biological basis that may underlie their unique vulnerability to the disruptive effects of obesity and the consumption of diets rich in saturated fats and sugars. Paralleling clinical data in humans (Geha et al., 2017; Riederer et al., 2016), we showed that DIO rats exhibit significant and partly irreversible microstructural alterations in the mPFC and amygdala regions associated with fear learning and fear extinction (Vega-Torres et al., 2018). The impact of obesogenic diets on PFC and BLA neuroplasticity is supported by studies showing reduced dendritic spine density in the PFC (Dingess et al., 2017) and dendritic length in the basal arbors of the BLA (Janthakhin et al., 2017) in rats that consume obesogenic diets rich in fats. Together, these structural alterations may lead to impairments in fear processing (Poulos et al., 2009).

It is becoming increasingly clear that alterations in reward-related brain areas play a major role in stress responsivity, with important implications for PTSD (Abraham et al., 2014; Corral-Frias et al., 2013; Holly and Miczek, 2016). Dopaminergic projections from the ventral tegmental area (VTA) to various forebrain areas including the accumbens (NAc) as well as the amygdala, PFC, hippocampus and hypothalamus participate in the reinforcing and motivational effects of several salient stimuli, regardless of their reward value. A growing body of evidence demonstrate that this dopamine system plays critical roles in fear learning and extinction in both humans (Haaker et al., 2013; 2015) and rodents (Fadok et al., 2009; Hitora-Imamura et al., 2015; Ng et al., 2018; Pignatelli et al., 2017; Zbukvic et al., 2017).

Several lines of evidence show that dopamine D1 receptors expressed in the NAc, the PFC, hippocampus and the amygdala play a critical role in fear learning and fear extinction (Abraham et al., 2016a; 2016b; Borowski and Kokkinidis, 1998; Fadok et al., 2009; Hikind and Maroun, 2008; J. J. Li et al., 2018; Ng et al., 2018). For example, mice lacking D1R exhibit deficits in both fear learning (Fadok et al., 2009) fear extinction (El-Ghundi et al., 2001). Further, studies show that systemic and intraamygdalar blockade of the D1 receptor with SCH-23390 attenuates FPS responses (Greba and Kokkinidis, 2000) and impair fear extinction in rats (Ng et al., 2018). There is some evidence that SCH23390 administration into the infralimbic region of the PFC but not the BLA suppresses the consolidation of fear extinction (Hikind and Maroun, 2008).

Work in both human and animals shows that obesity and intake of palatable and obesogenic high-fat diets leads to dysregulation in dopamine function. Please refer to de Macedo et al., and Reichelt for excellent reviews on this topic, (de Macedo et al., 2016; Reichelt, 2016). While initial intake of palatable and obesogenic diets increase dopamine levels, studies show a significant decrease in dopaminergic tone when animals acquire an obesogenic phenotype (J. Carlin et al., 2013; Norgren et al., 2006). Given the biphasic impact of diet on the dopaminergic system, it is reasonable that acute consumption of palatable foods will lead to a hyperdopaminergic state that may impair fear extinction; while chronic WD consumption will impair fear learning. The biology and phenotypes presented in this study strongly resembles those of a (mal)adaptive dopaminergic system.

In addition to changes in genes associated with dopamine uptake, metabolism, reduced D1 and D2 receptor expression levels have been documented in obesity (Alsiö et al., 2010; J. L. Carlin et al., 2016; L. M. Davis et al., 2009; Huang et al., 2005; Johnson and Kenny, 2010; Tomasi et al., 2015; Wang et al., 2001). Consistent with these studies and in particular Carlin et al., report (J. L. Carlin et al., 2016), our findings show reduced D1R mRNA levels in the PFC of the rats that consumed the WD during adolescence. The aforementioned studies demonstrate that high-fat diets, obesity, and fear conditioning alter dopamine receptor mRNA levels in the brain. Therefore, it is likely that a combination of these factors contributed to the observed differences in mRNA levels. Future studies are required to clarify the relative contribution of diet and fear conditioning to dopamine receptor gene expression. Given the critical role of PFC D1R in fear extinction, our findings suggest that diet-induced neuroadaptations in the dopamine system could have implications for stress-related psychopathology and PTSD.

### Limitations and Future Studies

This follow-up study had some limitations to be addressed in future research. The results should be interpreted with caution, as the study results of this rat model do not necessarily directly translate to the human condition. Although a key strength of longitudinal studies is the ability to measure change at the individual level and determine patterns of change across time, it is not clear from our data whether the impact of the WD on startle responses is a synergistic effect of brain maturation during adolescence or a diet effect. While studies suggest that adolescent rats are more vulnerable to the impact of obesogenic diets on aversive memories and PFC dopamine receptor 1 expression reduction (Boitard et al., 2015; J. L. Carlin et al., 2016), longitudinal studies with adult-onset WD intake are required to determine whether obesogenic diets synergize with brain development to alter fear responses or whether the diet alone can alter fear extinction correlates once the fear circuitry is fully mature. Whereas our results are consistent with deficits in fear extinction, it is unclear from our data whether the WD effects are related to impairments in fear extinction memory acquisition, consolidation, expression, reconsolidation and/or retrieval. Continued translational work will inform the basis of these learning and memory deficits. Lastly, replication in different rat strains, gender, and conditioning paradigms is warranted.

### Summary

This study shows that the consumption of obesogenic diets during adolescence heightens behavioral vulnerabilities associated with risk for anxiety and stress-related disorders. Given that fear extinction promotes resilience against PTSD and fear extinction principles are the foundation of psychological treatments for anxiety and stress-related disorders, understanding how obesity and obesogenic diets affect the acquisition and expression of fear extinction memories is of tremendous clinical relevance.

## Acknowledgements

This study was partly supported by the NIH (P20MD006988 and 2R25 GM060507) and the Loma Linda University School of Medicine Seed Grant Funds to JDF. We would like to thank the staff at the animal care facility.

## Financial Disclosures

All authors report no financial interests or potential conflicts of interest

